# Menstrual hygiene management practice and factors affecting it among high school females in Ambo City, Oromia state, Ethiopia, 2018: A cross-sectional mixed method

**DOI:** 10.1101/806422

**Authors:** Seifadin Ahmed Shallo, Wakeshi Willi, Abuzumeran Abubekar

**Author notes:** The corresponding author:, Institutional. this authors contributed equally to the work.

## Abstract

I.

**Background:** Though menstruation is normal physiology in the females’ life, if not managed safely, it can interrupt daily activity or it may lead to health problems. Poor water, sanitation and hygiene (WASH) facilities in schools, inadequate puberty education and lack of hygienic menstrual hygiene management items (absorbents) cause girls to experience menstruation as shameful and uncomfortable. This study aimed to assess the extent of unsafe menstrual hygiene practice and factors influencing it among school females in Ambo, Ethiopia.

**Methods:** an institutional-based mixed-method cross-sectional study was conducted from March 01 to 15/2018 to collect data from 364 school females, teachers, and investigators observation. To collect the data, self-administered technique, interview, observational and FGD techniques were used. Data were analyzed using SPSS statistical software version 20. Uni-variate, bivariate and multiple logistic regression analysis were done. With 95% CI, the P-value of less than 0.05 was taken as the level of significance.

**Results:** the prevalence of the unsafe menstrual hygiene management practice was 53.6%, which implies urgent response from the stakeholders is of paramount importance. Factors such as the age of the females, frequency of discussing menses with mothers and source of information about menses were variables significantly associated with menstrual hygiene management practice.

**Conclusion and Recommendation:** High numbers of school females’ menstrual hygiene management were poorly managed. This implies urgent measure is needed from the stakeholders to solve these problems so that sustainable development goal number 3, 4 and 5 will be achieved. To rid off the current problems which school females are facing, comprehensive and different sectors collaboration is important. Specifically, education sectors, water and sanitation sectors, and health sectors bear the frontline responsibilities.

## I. Introduction

### Background

Menstruation, a unique event in the life of developing females, is one of the milestones of puberty. It involves the cyclical shedding of the inner lining of the uterus which is controlled by the hormones produced by the hypothalamus and pituitary glands located in the brain(1).

Menstrual hygiene refers to personal hygiene practices during menstruation. This personal hygiene management starts with the choice of best sanitary materials, its proper utilization, disposal and body cleanliness(2). Menstruation is still regarded as something unclean or dirty in society. Seclusion of the menstruating girls/women and restrictions being imposed on them in the family not to participate in a different activity while on menses have reinforced a negative attitude towards this phenomenon(3).

Worldwide, nearly 52% of the female population (26% of the total population) is of reproductive age. The majority of these women and girls will menstruate each month for between two and seven days. Despite the challenges related to menstrual hygiene management, menstrual hygiene has been routinely ignored by professionals in the water sector, and in the health and education sectors too(4).

In Ethiopia, like in many parts of the African Countries, menstrual hygiene management is one of the critical problems adolescent girls facing while they are in school. This is mainly due to lack of infrastructure as well as poor management of the existing facilities which are important for the management of menstrual hygiene. Menstrual hygiene practices are clouded by taboos and socio-cultural restrictions even today, resulting from ignorance of the scientific facts and hygienic health practices, necessary for maintaining positive reproductive health(5).

Though menstruation is a naturally occurring event, it is associated with misconceptions, poor practices, and challenges among girls in developing countries. However, much is not documented; class-absenteeism and poor menstrual hygiene practices are common problems among adolescent girls.

In Ethiopia, 11% of girls change their menstrual cloths once a day. Girls commonly use rags torn from old clothes and use their dress tied in a knot to keep the sanitary cloths in place. Most girls rinse their rags in water without using soap; dry them under the bed; hang them in hidden, often unhealthy places in the house or on the roof, where they can grow mold. Such practices may increase susceptibility to infection(2).

Poor menstrual hygiene and inadequate self-care are major determinants of morbidity and other complications among the younger age group. Some of these problems include reproductive tract infections, urinary tract infections, scabies in the vaginal area, abdominal pain, absence from school, and complications during pregnancy(6).

In spite of these issues, menstrual hygiene has been largely neglected by the sector and other sectors focusing on sexual and reproductive health, and education. Even though the specific infections, the strength of the effect, and the route of transmission, remain unclear, it is plausible that poor MHM can affect the reproductive tract. Studies revealed that women who frequently use the reusable type of pad, and less frequently wash their genital during menses were at risk for lower reproductive tract infection, especially for bacterial vaginosis. Also, Baislye et. al reported that using cloths or cotton wool for menstrual hygiene is also a single most predictive for bacterial vaginosis infection(6)(7).

In a study conducted in Uganda among school girls, Julia et.al reported that about 90% of the schoolgirls didn’t meet the minimal criteria for safe MHM, which indicates the issue needs urgent responses from the concerned body. On the other way, the rate of genital irritation, discharge, and concern of malodor is higher among poor MHM girls. In addition to resulting in missing school, poor MHM also leads to a feeling of shame, worrying that odor may disturb the class and low self-esteem, which may results in psychological problems like depression during menses(8).

In the absence of parental and school support, girls cope, sometimes alone, with menses in practical and sometimes hazardous ways. Due to economic constraints or because of the females thought that it is taboo to ask parents for money to buy sanitary pads, they turn to transactional sex to get money for buying sanitary pads(9).

Menstruation is a silent issue in girl’s life which is further affected by the teacher’s perception and school resources. Because of this, many girls remain absent from schools during menstruation and sex education is also often neglected from the school curriculum which negatively impacts the student’s life. Educational policy analysis in lower and middle-income countries indicated that out of the 20 countries’ educational policy included in the study, the MHM didn’t get focus and even not mentioned in the all educational policy and in many of the countries’ school curriculum(10). In most of the schools, both male and female teachers are not ready to discuss menstruation and menstrual hygiene management with students. Many of the schools are male dominant and female teachers are not available. Teachers often skip such topics in books as they do not want any open discussion in the class or to escape from the questions asked by students. Also many were reported being absent from school due to lack of disposal system, broken lock/doors of toilets, lack of water tap, bucket, and poor water supply (11)(12).

The type of absorbents used during menses depends on the awareness of the existing materials, economic status, cultural acceptability, availability in the local market and personal preferences. Ideally recommended sanitary material for safe MHM is commercially available sanitary pads. However, though female reports they were using pads just not to be stigmatized, the most commonly mentioned alternatives were old clothes (rags), blanket or pieces of (bedding) mattress. Panties, socks, towel, cotton wool or tissue paper were also sometimes reported as absorbents for menses blood. When girls’ menses start unexpectedly, grass or leaves plucked from the ground around the schoolyard was reported to be the only option that schoolgirls have once they faced menses at school(9).

Proper disposal of used menstrual material is still a challenge for many countries of the world. Because of this, most of the women dispose of their sanitary pads or other menstrual articles into household solid wastes or garbage bins that ultimately become a part of solid waste. In countryside areas, most women use reusable and non-commercial sanitary materials like reusable pads or cloths. Thus, they generate a lesser amount of menstrual waste as compared to women in urban areas who rely on commercial disposable pads(13). While they are in the schools, due to the lack of hygienic facilities like dust bins, girls throw their pads in the toilets. In some cases, girls threw away their used menstrual clothes without washing them. As sanitation systems were designed with urine and feces in mind, they are unable to cope with the menstrual absorption materials and will block the sewerage and causes blockage or backflow(12).

In general, different studies were conducted on the issue related to menstruation and menstrual hygiene management among school females in Ethiopia. However, the way they measured the hygienic practice of menstrual hygiene varies. Many of the studies use practice assessing questions and based on the number of items they grouped MHM as good or bad or as poor or good. We believe that such classification is not objective, not comprehensive and it doesn’t meet with the actual/standard definition of menstrual hygiene management. Because of this, it is even difficult to compare or to conclude the extent of poor menstrual hygiene management with the currently existing works of literature in the country.

So, this study assessed practices of menstrual hygiene management and associated factors among high school Females in Ambo Town West Showa Zone, Ethiopia, using the four major areas of the MHM which is in line with the internationally recognized definition of MHM with the following specific objectives.
1. What extents of high school females were practicing unsafe MHM?
2. What are the underlying determinants of MHM practices among high school Females?

## II. METHODS

### Study area

The study was conducted in Ambo town, Oromia regional state, Ethiopia. With a population of about fifty thousand, Ambo is located 112 km west of Addis Ababa. Ambo is the capital city of West Shoa Zone. Concerning the education facility, there was one government & one private University, three private colleges, two Preparatory schools, five high schools and seventeen primary schools in the Ambo Woreda.

### Study Design and Period

The institutional-based cross-sectional study design was conducted and data was collected from March 01 to 15/2018.

### Source population

The source populations were all female students attending high schools in Ambo city, West Shoa Zone, Oromia regional state.

### Inclusion and exclusion criteria

All female students who started menses were included in the study.

### Sample size and Sampling Techniques

There were five high schools (4 governmental and 1 private) in Ambo town. In the city, the total number of students enrolled in high school levels for the academic year 2017/18were 5230 out of which 2829 were males and 2401 were females.

The sample size was determined based on the assumption of simple random sampling (SRS) method using the formula for a single population proportion. Since we were unable to get a proportion from the previous study which is similar to our study plan, we decided to take P=50% assuming that fifty percent of school females were practicing unsafe MHM. Using 0.05 margin of error, the sample size was calculated as follows:

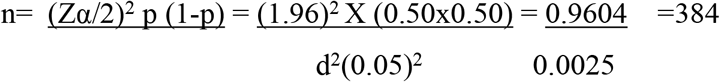

**Where,**

n= the desired sample size

p= Prevalence of schoolgirls who expected to practice poor MHM

Z= is the standard normal score set at 1.96 (95% confidence interval)

d= is the margin of error to be tolerated (5%)

Since the source population was finite, the sample size correction approach was applied and a 10% nonresponse rate was added. The final sample size was 364.

### Sampling procedure

There were five high schools in Ambo town (4 governmental and 1 private). They were: Ambo secondary school, Bakalcha Secondary school, Awaro Secondary school, Liben Macha secondary school & Future generation hope secondary School. The participants from each School were selected using probability proportional to the number of female students they had. The schematic procedure for selecting the study participants is described in figure elsewhere. (Fig 1).

**Fig. 1.**
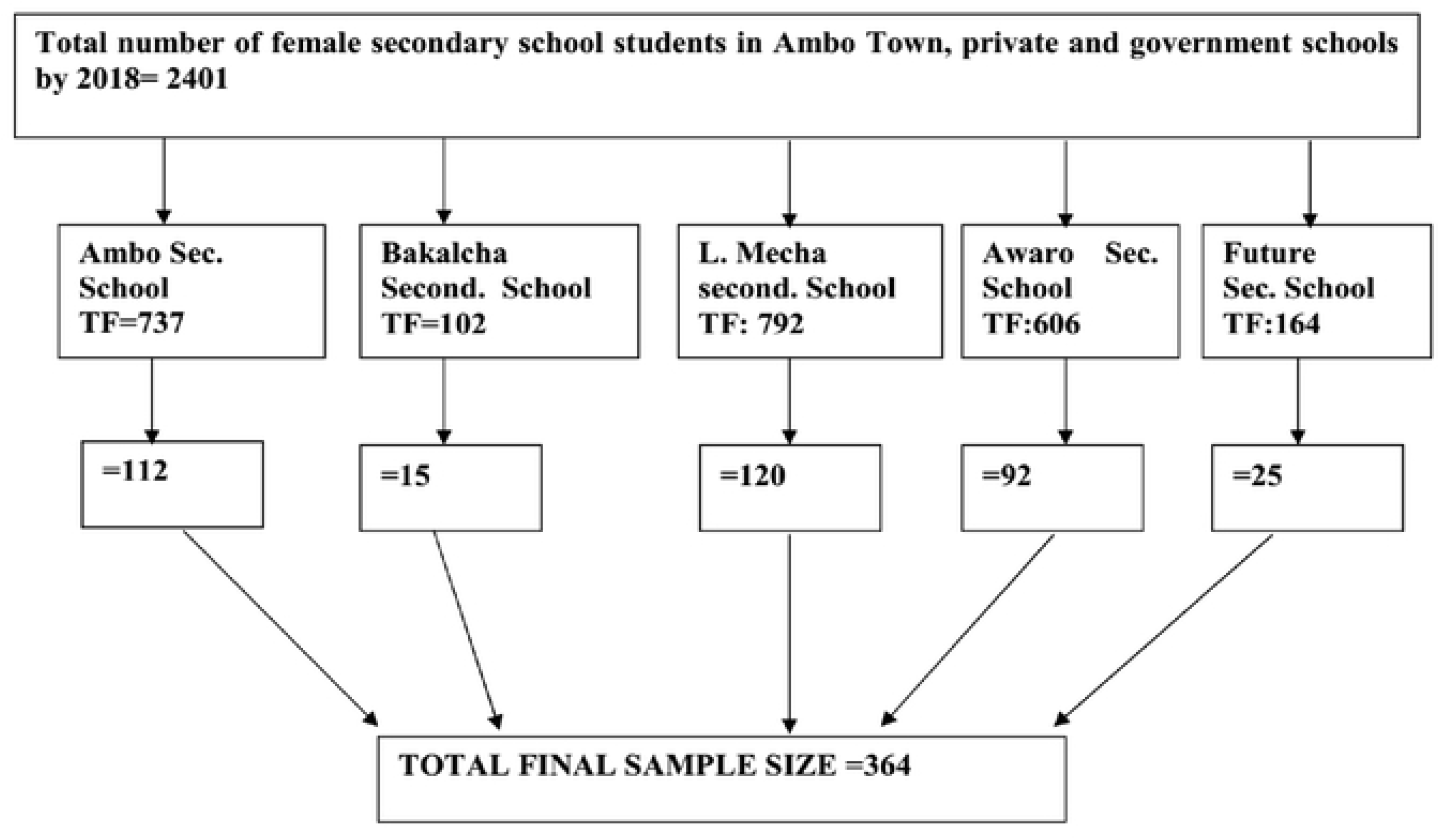
Schematic diagram presenting the sampling procedure from five high schools in Ambo city, Oromia, Ethiopia, 2018.

### Data collection tools and methods

To collect the data, self-administered technique, interview, observational and FGD techniques were used. Quantitative data were collected using structured and pretested questionnaire containing both open-ended and closed-ended questions. The questionnaire was adopted from different works of literature developed for similar purposes by different authors and then modified to suit the local condition so that it doesn’t contradict with the local cultures and beliefs. Five female data collectors, diploma holders, were recruited and trained for two days on the objective of the study, how to support participants and how of consent taking from participants. The questionnaire was prepared first in English and then translated into Afan Oromo and Amharic (the commonest local languages).

For qualitative data, a semi-structured interview guide was prepared to collect data from school director. Observational checklist which assesses the presence of separate toilets for males and females, presence of water around the toilet, cleanness of the toilet, presence of waste disposal bins in /around the toilet and other related issues were prepared and utilized.

Five focus group discussions (FGDs) (one at each school) were conducted to gather information from a total of 36 school females aged 15 to 20. The females were selected purposively by the help of the school director based on their school performance, age and their residence so that the group will be balanced in its structure. The FGDs and in-depth interview was tape-recorded and transcribed later. Data analysis was done through a thematic analysis approach. The patterns of experiences were derived from the transcripts, either from direct quotes or through paraphrasing common ideas. Data from all the transcripts relating to the classified patterns were identified and placed under the relevant theme where it complements the quantitative findings.

### Data quality assurance

To ensure the quality of the collected data, data collectors were trained well, the principal investigators and filled supervisors check the completeness and clarity of the questionnaire immediately after received from the respondents in the field.

### Data management and analysis

The collected data were checked visually for completeness then, coded and entered into Epi data version 4.5 statistical packages. Descriptive analysis such as frequency distribution with percentage was computed. To assess the association between dependent and independent variables by controlling for confounders, first binary logistic regression was run and variables with p-value <=0.25 and the variables which are known to have an association with dependent variables from reviewed literature were selected for Multiple logistic regression analysis. Statistical significance was declared at P-value <0.05 with 95% confidence interval (CI).

### The study variables

#### Dependent Variables

Menstrual hygiene management practice (safe or unsafe)

#### Independent Variables

Age of respondents, Place of residence, Religion, living condition, sexual activity, accessibility of constant pocket money, parental factors like educational status, Educational level of parents, Accessibility of sanitation facilities in school compounds, etc.

### Operational definitions

#### Safe Menstrual Hygiene practice

the menstrual hygiene practice is considered to be safe if it fulfills all of the following four criteria, otherwise unsafe. If the females used safe absorbents (considered safe if they were commercially available sanitary pads (locally called modes/TAMPON) or new clothes). Old clothes and other items such as toilet paper, mattress, sponge or underwear alone considered unsafe. Changing absorbents three or more times per 24 hours, If girls wash their genitalia two or more times per day and disposed of used menstruation pad by burying or if burnt it after use(8)(14).

### Ethical considerations

After the proposal passed through different stages, Ethical clearance was obtained from the ethical review committee of the College of Medicine and Health Sciences, Ambo University. During the fieldwork, the objective of the study was clearly explained for the study participants, the confidentiality of the data to be collected and the right not to participate was also assured. Consent forms were distributed for each of the participants who were age greater than or equal to 18 years and after they read the consent form, the participants were asked to confirm their participation in the study by signing the consent form. For those who were aged less than 18 years, an assent letter was sent to their family through the students. Accordingly, all of the Females family signed and sent it back the assent for us.

We used the STROBE cross-sectional reporting guidelines(15).

## III. Results

Out of the 364 participants intended to be included, 336(91.5%) were responded fully to the distributed questionnaire. The mean ages of the participants were 16± 2.25 SD years. The majority of the students were urban residents (92.9%), single in marital status (93.8%), at the age of less than or equal to 17 years (92.9%) (Table-1).

**Table 1:** socio-demographic distribution of in school Females in Ambo City, 2018.

| Variables | Frequency | Percent (%) |
| --- | --- | --- |
| **Age** |  |  |
| 12-17 | 312 | 92.9 |
| >18 | 24 | 7.1 |
| **Residence** |  |  |
| Urban | 312 | 92.9 |
| Rural | 24 | 7.1 |
| **Marital status(relationship)** |  |  |
| single | 315 | 93.8 |
| Engaged | 21 | 6.2 |
| **Religion** |  |  |
| Orthodox | 158 | 47.0 |
| Protestant | 142 | 42.3 |
| Wakefata | 36 | 10.7 |
| **Mothers Educational status** |  |  |
| illiterate | 66 | 19.6 |
| 1-6 | 103 | 30.7 |
| 7-12 | 83 | 24.7 |
| Degree and above | 81 | 24.1 |
| **Father's Educational status** |  |  |
| Illiterate | 59 | 17.6 |
| grade 1-6 | 58 | 17.3 |
| 7-12 | 111 | 33.0 |
| Degree and above | 107 | 31.8 |
| **With whom do you live** |  |  |
| With both parents | 211 | 62.2 |
| only with mother | 80 | 23.6 |
| Alone | 13 | 3.8 |
| with relatives | 32 | 9.4 |

### Menstrual Hygiene management related Awareness and practice of the respondents

In this study, the mean age at menarche was 13.9± (0.71SD). About 175(51.7%) of the respondents reported that their menses stayed up to seven days per cycle. The majority (about 73%) reported that they had awareness about menses before menarche. Mothers were the leading (47%) source of information about menses for the respondents. About 82.2% of the females reported that it is easy for them to discuss menses with their mothers. The respondents were asked “how you rate your preparedness for the first menses?” with the option of not at all, well prepared, not well prepared or I don’t remember. Accordingly, only 31% reported that they prepared well for the first menses while the others reported not prepared well or don’t remember. About 81(24%) of the participants don’t know the existence of commercially available sanitary pads. Nearly 17% of the respondents reported that the source of menses blood is organs other than Uterus like abdomen, vagina and or others. The respondents were also asked “what is the cause for the menses bleeding?” and about 26% reported it is God curse or disease (Table-2).

**Table: 2.** Menstrual Hygiene management related Knowledge among high school females in Ambo City, Ethiopia, 2018.

| **Variables** | **Frequency** | **Percent** |
| --- | --- | --- |
| **Awareness about menses before menarche** |  |  |
| Yes | 247 | 72.9 |
| No | 88 | 26.0 |
| **The main source of information about Menses** |  |  |
| mother | 157 | 46.3 |
| teacher | 96 | 28.3 |
| friends | 57 | 16.8 |
| Others(books, media) | 23 | 6.8 |
| **What is the cause for menses** |  |  |
| hormone | 235 | 69.3 |
| curse of God | 80 | 23.6 |
| caused by disease | 16 | 4.7 |
| **How much you prepared for the first menses** |  |  |
| not at all | 136 | 40.1 |
| not well prepared | 23 | 6.8 |
| prepared well | 100 | 29.5 |
| I don't remember | 73 | 21.5 |
| **From where does menses blood come** |  |  |
| uterus | 271 | 79.9 |
| Other body organs (vagina, bladder, abdomen) | 58 | 17.1 |
| **It is harmful to exercise during menses** |  |  |
| Yes | 185 | 54.6 |
| No | 141 | 41.6 |
| **Poor MHM exposes for infection** |  |  |
| yes | 295 | 88.1 |
| no | 40 | 11.9 |
| **Menses Related school absenteeism in the last 3 months** |  |  |
| Yes | 108 | 32.1 |
| No | 228 | 67.9 |
| **Why you absent from school during menses\*\*** |  |  |
| I can't access sanitary pad | 59 | 54.6 |
| No suitable place for pad exchange | 67 | 62 |

\*\* Since multiple responses are possible, the percent could be greater than 100%

Different works of literature used different criteria to classify menstrual hygiene management as good or poor, or as safe or unsafe. For instance, many studies in different parts of Ethiopia and African Countries used item questions related to practice and the mean score was taken as the cutoff point for classifying MHM status(16)(17).

Out of the total 336 study participants, nearly 25% of them reported that they don’t know the presence of commercially available sanitary pads and 12% reported that they used rag clothes for managing their menses blood in the recent menses. Ten percent of the respondents reported that they disposed of the sanitary pad/materials they used in open filed. The respondents were also asked how could they managed their menses if faced at school and about 32% reacted that they leave home if faced suddenly bleeding.

Concerning the frequency of cleaning the genitalia during menses, 43.7% reported that they have cleaned the genital area with soap and 16.5% cleaned their genitalia with tissue paper or piece of clothes.

In order to come up with the comprehensive MHM practice as safe or unsafe, we used the four major criteria from UNICEF definition of menstrual hygiene i.e. MHM is Women and adolescent girls use clean material to absorb or collect menstrual blood, and this material can be changed in privacy as often as necessary for the duration of the menstrual period. MHM includes using soap and water for washing the body as required and having access to facilities to dispose of used menstrual management materials (2). In this study, menstrual hygiene management is considered to be safe if it fulfills the following criteria otherwise considered being unsafe. If the females used safe absorbents (considered safe if they were commercially available sanitary pads (locally called modes) or new clothes. Old clothes and other items such as toilet paper, mattress, sponge or underwear alone considered unsafe. Changing absorbents three or more times per 24 hours, If girls wash their genitalia two or more times per day and disposed of used menstruation pad by burying, in the toilet or if burnt it after use.

Using these criteria, Out of the total 336 respondents, about 53.6% (95% CI: 48.5, 58.6) of them practiced unsafe menstrual hygiene management. Of the four items used for grading the menstrual hygiene management, the most poorly managed was the frequency of washing genitalia in which 28% of the female didn’t wash their genitalia during bleeding till blood stop and 48.7% reported they clean their genitalia every two days or more (table-3).

**Table: 3.** Menstrual Hygiene practice among School females in Ambo city, 201.

| **Variables** | **Frequency** | **Percent** |
| --- | --- | --- |
| **Do you know the commercially available sanitary pad** |  |  |
| yes | 255 | 75.1 |
| no | 81 | 24.9 |
| **Type of sanitary pad used mostly** |  |  |
| Rag (cloth of any kind) | 42 | 12.4 |
| piece of clean clothes | 63 | 18.6 |
| disposable sanitary pads | 218 | 64.3 |
| pant alone | 10 | 2.9 |
| **Frequency Of sanitary pad change** |  |  |
| once per day | 53 | 15.6 |
| twice per day | 245 | 72.3 |
| three times and above | 37 | 10.9 |
| **How do you manage Menses at School(if faced)** |  |  |
| used modes carry in pocket school | 180 | 53.1 |
| left school | 111 | 32.7 |
| do nothing | 41 | 12.1 |
| **Disposing sanitary pad** |  |  |
| In toilet | 228 | 67.3 |
| In dust bin | 71 | 20.9 |
| open field | 34 | 10.1 |
| **how do you clean your body during menstruation** |  |  |
| take bath with soap | 129 | 38.1 |
| clean around genitalia with tissue paper/piece of clothes | 56 | 16.5 |
| wash genitalia area with soap | 148 | 43.7 |
| **Frequency of cleaning genitalia during Menses** |  |  |
| daily | 70 | 20.6 |
| every two day | 165 | 48.7 |
| do not take a bath at all while on the bleeding | 95 | 28.0 |
| **Do you usually get pocket money for Sanitary Pad** |  |  |
| Yes | 255 | 75.9 |
| No | 81 | 24.1 |
| **Overall MHM status** |  |  |
| **Safe \*** | 157 | 46.7 |
| **Unsafe** | 179 | 53.3 |

The data collected through observation indicated that all schools (4 both public and one private) have latrines in the school compound. Each toilet is separated and labeled for both men and females. But, except one which is a private school, the others have poorly maintained. They have no water source or nonfunctional water source, no door, and locks. The toilets were not clean enough; even some of them were filled, and difficult to utilize. In general, it lacks proper waste disposal, hand washing and menstrual hygiene facilities which makes it difficult for girls to change and dispose of absorbent materials secretly.

### Factors Associated with unsafe menstrual hygiene management

Both bivariate and multivariable logistic regression analyses were done to identify factors associated with MHM practices. In bivariable analysis: age, marital status, religion, parental educational status, respondents living status (with whom she is living currently), the major source of information about menses, frequency of discussing menses with their mother have an association with unsafe MHM practice. After running multiple variable analyses, variables like age, religion, father educational status, frequency of discussing menses with their mother were significantly associated with unsafe menstrual hygiene management practice.

Accordingly, compared to age greater or equal to 18 years, those females in the age of less than 18 years were 84% less likely to be their MHM practice is unsafe 95% CI, [AOR: 0.16(0.045, 0.57), P=0.005]. Females whose fathers were attended degree and above were 72% with [AOR: 0.28, 95% CI: 0.10, 0.88, P-value of 0.03] less likely their menses is unsafely managed compared to those whose father can’t read or write. Compared to those female who reported that they almost never discuss about menses with their mother, females who reported they frequently discuss about menses with their mother (think that it’s easy or very easy for them to talk about menses with their mother) were 70%, [AOR: 0.30, 95% CI: 0.13, 0.71, P-value of 0.006] less likely their menstrual hygiene management practice was unsafe.

Compared to those females who reported that they used media (electronic/books) as common source of information about menses related issue, those who claimed their school teacher were common source of info on menstrual issue were 3.75 times more likely to be unsafe (AOR:3.75, 95% CI: 1.75, 8.00) P= 0.001). (Table-4)

**Table: 4.**
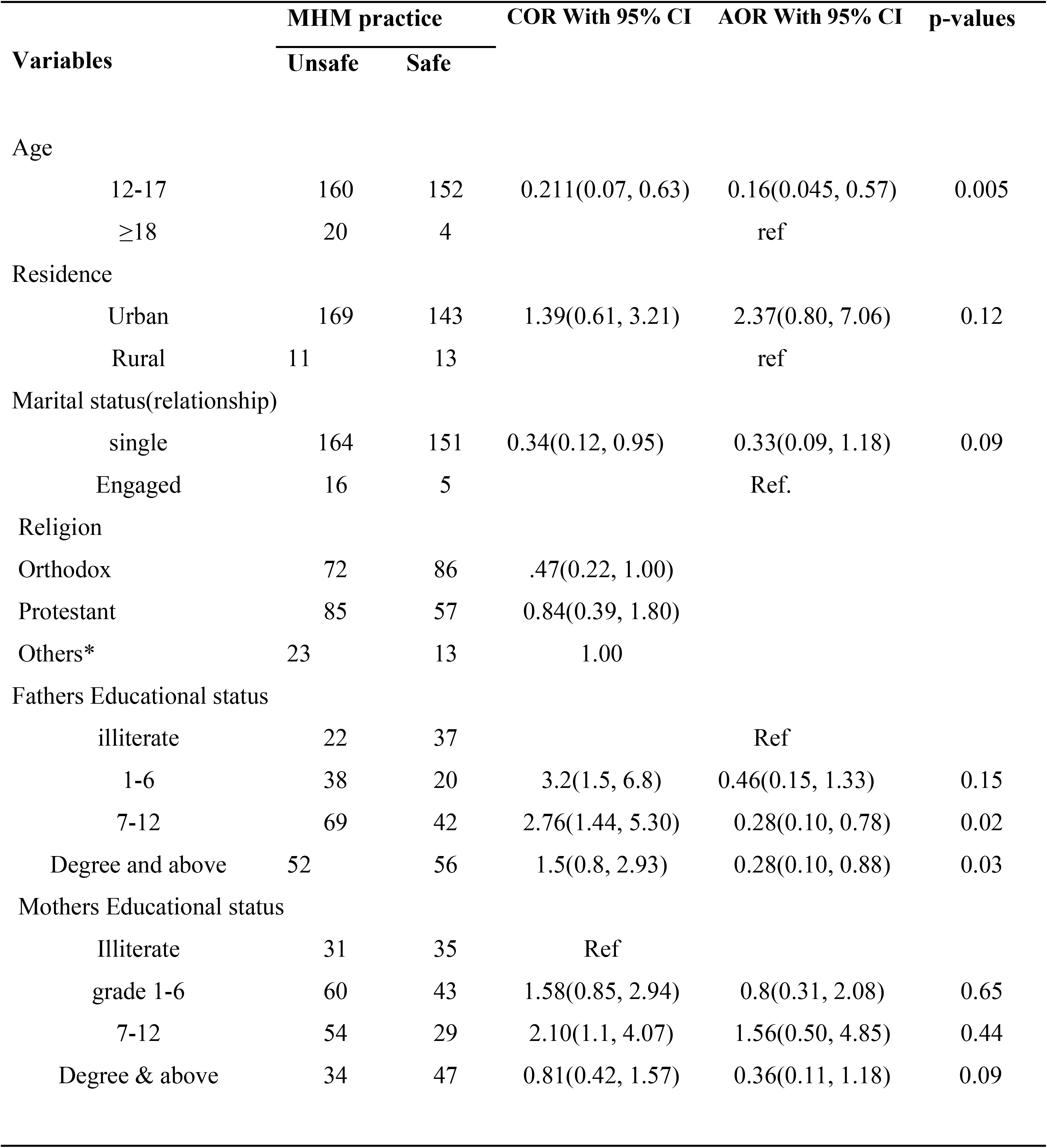
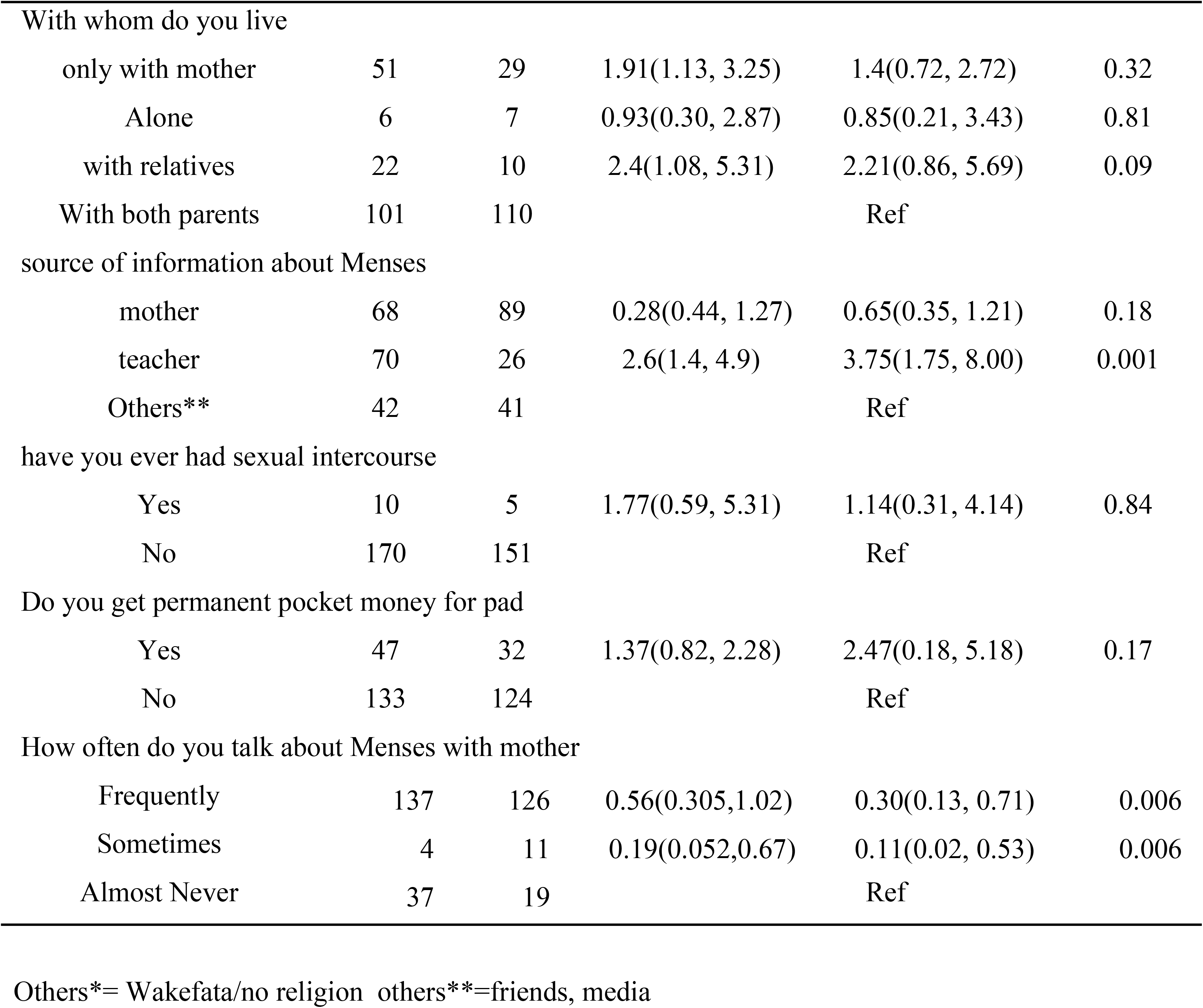
Bivariate and Multivariable logistic regression analysis for factors associated with unsafe menstrual hygiene management among high school females in Ambo city, West Shoa, Oromia regional state, Ethiopia, 2017.

| Variables | MHM practice |  | COR With 95% CI | AOR With 95% CI | p-values |
| --- | --- | --- | --- | --- | --- |
|  | Unsafe | Safe |  |  |  |
| Age |  |  |  |  |  |
| 12-17 | 160 | 152 | 0.211(0.07, 0.63) | 0.16(0.045, 0.57) | 0.005 |
| ≥18 | 20 | 4 |  | ref |  |
| Residence |  |  |  |  |  |
| Urban | 169 | 143 | 1.39(0.61, 3.21) | 2.37(0.80, 7.06) | 0.12 |
| Rural | 11 | 13 |  | ref |  |
| Marital status(relationship) |  |  |  |  |  |
| single | 164 | 151 | 0.34(0.12, 0.95) | 0.33(0.09, 1.18) | 0.09 |
| Engaged | 16 | 5 |  | Ref. |  |
| Religion |  |  |  |  |  |
| Orthodox | 72 | 86 | .47(0.22, 1.00) |  |  |
| Protestant | 85 | 57 | 0.84(0.39, 1.80) |  |  |
| Others* | 23 | 13 | 1.00 |  |  |
| Fathers Educational status |  |  |  |  |  |
| illiterate | 22 | 37 |  | Ref |  |
| 1-6 | 38 | 20 | 3.2(1.5, 6.8) | 0.46(0.15, 1.33) | 0.15 |
| 7-12 | 69 | 42 | 2.76(1.44, 5.30) | 0.28(0.10, 0.78) | 0.02 |
| Degree and above | 52 | 56 | 1.5(0.8, 2.93) | 0.28(0.10, 0.88) | 0.03 |
| Mothers Educational status |  |  |  |  |  |
| Illiterate | 31 | 35 | Ref |  |  |
| grade 1-6 | 60 | 43 | 1.58(0.85, 2.94) | 0.8(0.31, 2.08) | 0.65 |
| 7-12 | 54 | 29 | 2.10(1.1, 4.07) | 1.56(0.50, 4.85) | 0.44 |
| Degree & above | 34 | 47 | 0.81(0.42, 1.57) | 0.36(0.11, 1.18) | 0.09 |
| With whom do you live |  |  |  |  |  |
| only with mother | 51 | 29 | 1.91(1.13, 3.25) | 1.4(0.72, 2.72) | 0.32 |
| Alone | 6 | 7 | 0.93(0.30, 2.87) | 0.85(0.21, 3.43) | 0.81 |
| with relatives | 22 | 10 | 2.4(1.08, 5.31) | 2.21(0.86, 5.69) | 0.09 |
| With both parents | 101 | 110 |  | Ref |  |
| source of information about Menses |  |  |  |  |  |
| mother | 68 | 89 | 0.28(0.44, 1.27) | 0.65(0.35, 1.21) | 0.18 |
| teacher | 70 | 26 | 2.6(1.4, 4.9) | 3.75(1.75, 8.00) | 0.001 |
| Others** | 42 | 41 |  | Ref |  |
| have you ever had sexual intercourse |  |  |  |  |  |
| Yes | 10 | 5 | 1.77(0.59, 5.31) | 1.14(0.31, 4.14) | 0.84 |
| No | 170 | 151 |  | Ref |  |
| Do you get permanent pocket money for pad |  |  |  |  |  |
| Yes | 47 | 32 | 1.37(0.82, 2.28) | 2.47(0.18, 5.18) | 0.17 |
| No | 133 | 124 |  | Ref |  |
| How often do you talk about Menses with mother |  |  |  |  |  |
| Frequently | 137 | 126 | 0.56(0.305,1.02) | 0.30(0.13, 0.71) | 0.006 |
| Sometimes | 4 | 11 | 0.19(0.052,0.67) | 0.11(0.02, 0.53) | 0.006 |
| Almost Never | 37 | 19 |  | Ref |  |

Others\*= Wakefata/no religion others\*\*=friends, media

## IV. Discussion

In this study, about 53.6 %(95% CI: 48.5, 58.6) of the school females MHM practice was unsafe. This finding is lower when compared to studies conducted in Bahir Dar, in Nekemte, Ethiopia, and in Uganda in 2014/15 among school girls, in which the prevalence of unsafe MHM practice was 75.5%, 60%, and 90.5% respectively(7,13,14). The observed difference in prevalence could be due to the difference in the year of the study period. The two previous studies were conducted almost four years back. Over this long period, there might be an improvement in accessibility and availability of sanitary pad, improvement in the awareness of females about menstrual hygiene, improvements in water and sanitary (WASH) facilities in the school environment. For instance, about 64.3% of respondents used a sanitary pad in the current study and this is higher compared to the previous study conducted in Bahir DDar and Wollo in which only 44.9 % and 33% reported using sanitary pads during the recent menses respectively(18)(14).

The majority (46.3%) of the study participants reported that the main source of information about menses was their mother, which is similar to a study conducted in Nekemte, west Oromia and in sokota, Nigeria(14, 17). But, it is, in contrast, to study in the Northern part of Ethiopia by Population council office in which the majority (40%) of females received information about menses from their teacher instead of mothers(9%) (17). This difference is most probably because of cultural differences. In the Northern part of Ethiopian regions, where child marriage is high, females/girls are considered as ready for marriage or a sexual debut if she started experiencing menses. To escape this, females may hide their menses from their mothers and request teachers for information about menses(20).

Even though the school females were reporting they get information from different sources like parents, teachers, friends, and media, they don’t believe that the information they were getting is sufficient enough in enabling them to practice safe MHM practice. The information they get is too minimal, they shy to ask detail about menses in the class and it is also taboo in the community to talk detail about menses with parents. Even what they commonly ask their teacher was asking permission to leave class for menses reason or describing menses as a reason for absent from class. Because of this, the FGD discussants urge if there is an accessible way of communicating detail information about menses and its safe management.

*“The majority of our school females lack enough information on how to manage their menstruation and afraid to ask or discuss the issue of the menstrual cycle with their teachers or parents. We didn’t get any readable material which prepared with the local language for better information and knowledge.”(18 years FGD discussants from Ambo secondary School)*

About 32% of school females reported that they left school if they suddenly faced menses at school. This is most probably because of the lack of facilities necessary for menstrual blood management. Our observational finding at school also indicated that there were toilets in all schools and separated for both males and females. But, some of them lack doors which are not conducive for privacy, almost all of them have no water facilities, and some of them were dirty.

“*The school compound and latrine area has no concerned cleaner, and because of unavailability of water, the toilet is always dirty and offensive which is not safe to use or change modes*” (female FGD discussant from Bakalcha school)

Lack of water and sanitary facility is also contributing to school absenteeism. For example, about 32% of the respondents were absent at least once in the last three menstrual cycles for a reason related to menses. Out of them, 54% reported that they were absent from the class because of unable to get a safe sanitary pad. The FGD finding also supports this finding.

“*Usually I don’t come to class during my period eve. Because, if the period comes suddenly while I am at school, my attention becomes drown toward fear of having an accidental leakage and thinking that if my cloth suddenly stained with leakage of blood the students may laugh at me, so that I preferred staying at home”* (16 years old FGD discussant from Liban Mecha school).

Even when female students come to school while they were on menses, their feeling is easily understandable by teachers as they are on the menses. One of the female school teachers portrayed the status of females on menses as following:

*“We know their conditions in the class if one female student is on her period. She is not attentive and looks worried, frequently changes her place to back and left the class after all students leave the class”* (A Female teacher from one of the school)

More than 32% of the females used homemade alternative sanitary materials for managing their menses blood. This homemade alternative material reaches from old clothes/rags to clean new clothes. These old clothes may put the women at risk for infection and allergic reactions for their skin around the genitals. This finding is similar to the study conducted in Udaypur, Nepal(21).

The MHM practice was varied by females’ age. Those females in the age of less than 18 were 84% [95%, AOR: 0.16, CI: 0.05, 0.57] less likely their menstrual hygiene management practice was unsafe. This could be because most of the time younger age groups are more likely to be supported by family compared to their older sisters and older age females are considered as they are already aware of about menses and related issues(20).

The other variables related to MHM status is the frequency of discussing menses with parents. In this study, compared to females who rarely talk about menses with their mothers, those who frequently discuss with mothers were 70% less likely [95% AOR: **0**.30 CI: 0.13, 0.71] their MHM is unsafe. This is because the more the female discuss menses with their family about menses and another SRH issue, the more likely they will develop confidence and get support from their parents and even easy for them to request for money to buy a sanitary pad, soap and other materials for sanitation.

### Limitation of the study

The menstrual hygiene issue is something considered taboo in the community and too sensitive to discuss openly. This may contribute to social desirability bias. So the interpretation of this finding should put this into consideration. In addition, since this is a cross-sectional study, it doesn’t necessarily indicate cause and effect relationships.

## Conclusion and Recommendation

Generally, we have tried to estimate the prevalence of unsafe menstrual hygiene management with the concept definition of MHM and its associated factors. Accordingly, we can conclude the following from the above finding:

- A high number (53.6%) of school females’ MHM is unsafe. This poor management of MHM practice may put them at risk for reproductive organ infection, allergic reaction around the genital area, fear of smell, shyness, low self stem finally may be ended up with school abstention and low school performance.
- About 32% of school females absent from school due to menses related reason especially water, sanitation, and hygiene facility-related problems in the schools. This gap is one of the factors which may affect the achievement of sustainable development goal number 4, 5, and number 6 directly.
- Almost all of the schools were not female friendly for managing their menstrual hygiene.
- Female students urge for detail information about menses, MHM, and affordable sanitary materials.
- Age of the females, frequency of discussion with their parents, source of information and fathers educational status were significantly associated with MHM status

Based on the above finding, we would like to recommend the following:

To rid off the current MHM challenges that school females are facing, comprehensive and different sectors collaboration is important. Specifically, education sectors, WASH sectors, and health sectors bear the lion shares. To achieve safe MHM at school level:

Females should get detail information about menses, its safe management and the consequences of poorly managed menses blood. This can be achieved through continuous awareness creation by establishing clubs mainly working on MHM, regularly preparing and distributing pamphlets in the local language. For the achievements of the above things, the schools should work with its stakeholders including health sectors.

In addition, the educational office should urgently emphasize on fulfilling water, sanitation and hygiene facilities at school, and regular monitoring of those existing to be fully functional.

Government and nongovernment organizations working on the WASH sector should also give focus on availability, accessibility, and affordability of menstrual hygiene pads.

Any programs working to improve the MHM should target the age of the female, mothers and teachers, sustainable information source for females so that their program will be a success.

Finally, further studies are needed on the type of sanitary pad that that is preferable, socially acceptable, sustainable, and environmentally friendly.

## Lists of abbreviations and Acronyms

AOR: Adjusted odd ration
CI: confidence interval
FGD: focus group discussion
MHM: menstrual hygiene management
SD: standard deviation
WASH: water and sanitation hygiene

## Declaration

### Ethical approval and consent to participate

The proposal for the study was submitted to the Ethical Review Committee of College of Health and Medical Sciences, Ambo University for approval and clearance. Accordingly, the study was checked for its ethical issue and a permission letter was obtained. The letter for support was written from the college of medicine and Health Sciences, Ambo University to all selected Schools.

### Consents from participants

Before starting the data collection process, both written and informed verbal consent was taken from each respondent and their families. For those females age less than 18 years, their families were contacted and assent was taken after briefing about the objectives of the study.

### Consent for publication

Not applicable

### Availability of data and material

All data generated or analyzed during this study were included in this published article and its supplementary information files are in the hands of the correspondent author.

### Authors’ contributions

**SAS** developed the proposal, analyzed data, and the major contributor of the manuscript.

**WWM** had participated in data collection supervision, read the study throughout the progression of study, starting from proposal development to data analysis.

AA: has participated in the designing, data collection and manuscript preparation.

All authors read and approved the final manuscript.

## Acknowledgments

We would like to acknowledge Ambo University for covering the data collectors’ costs and supervisors per diem. For this study and we would like to thank our study participants, our data collectors for their collaboration.

## REFERENCES

1. Santra S. Assessment of knowledge regarding menstruation and practices related to maintenance of menstrual hygiene among the women of the reproductive age group in a slum of Kolkata, West Bengal, India. Int J Community Med Public Heal [Internet]. 2017 Feb 22 [cited 2019 Apr 28];4(3):708. Available from: http://www.ijcmph.com/index.php/ijcmph/article/view/1124

2. UNICEF and Emory University. WASH in Schools Empowers Girl’s Education. Tools for Assessing Menstrual Hygiene Management in Schools. UNICEF and Centre for Global Safe Water, Emory University: New York, 2013. [Internet]. 2013 [cited 2019 Apr 15]. Available from: www.unicef.org/wash/schools

3. Lawan UM, Yusuf NW, Musa AB. Menstruation and menstrual hygiene amongst adolescent school girls in Kano, Northwestern Nigeria. Afr J Reprod Health [Internet]. 2010 Sep [cited 2019 Apr 28];14(3):201–7. Available from: http://www.ncbi.nlm.nih.gov/pubmed/21495614

4. House S, Mahon T, Cavill S. Menstrual Hygiene Matters: a resource for improving menstrual hygiene around the world [Internet]. Vol. 21, Reproductive Health Matters. Taylor & Francis, Ltd.; [cited 2019 Jul 10]. p. 257–9. Available from: https://www.jstor.org/stable/43288983

5. Ayalew Tegegn+ Meseret Yazachew+ Yeshigeta Gelaw+. Reproductive Health Knowledge and Attitude among Adolescents: A community-based study in Jimma Town, Southwest Ethiopia | The Ethiopian Journal of Health Development (EJHD) [Internet]. [cited 2019 Apr 28]. Available from: https://www.ejhd.org/index.php/ejhd/article/view/508

6. Baisley K, Changalucha J, Weiss HA, Mugeye K, Everett D, Hambleton I, et al. Bacterial vaginosis in female facility workers in north-western Tanzania: prevalence and risk factors. Sex Transm Infect [Internet]. 2009 Sep 1 [cited 2019 Apr 28];85(5):370–5. Available from: http://www.ncbi.nlm.nih.gov/pubmed/19473997

7. Torondel B, Sinha S, Mohanty JR, Swain T, Sahoo P, Panda B, et al. Association between unhygienic menstrual management practices and prevalence of lower reproductive tract infections: a hospital-based cross-sectional study in Odisha, India. BMC Infect Dis [Internet]. 2018 Dec 21 [cited 2019 Apr 28];18(1):473. Available from: http://www.ncbi.nlm.nih.gov/pubmed/30241498

8. Hennegan J, Dolan C, Wu M, Scott L, Montgomery P. Measuring the prevalence and impact of poor menstrual hygiene management: a quantitative survey of schoolgirls in rural Uganda. BMJ Open [Internet]. 2016 Dec 30 [cited 2019 Apr 4];6(12):e012596. Available from: http://bmjopen.bmj.com/lookup/doi/10.1136/bmjopen-2016-012596

9. Mason L, Nyothach E, Alexander K, Odhiambo FO, Eleveld A, Vulule J, et al. ‘We Keep It Secret So No One Should Know’ – A Qualitative Study to Explore Young Schoolgirls Attitudes and Experiences with Menstruation in Rural Western Kenya. Molyneux C ‘Sassy’, editor. PLoS One [Internet]. 2013 Nov 14 [cited 2019 Apr 28];8(11):e79132. Available from: https://dx.plos.org/10.1371/journal.pone.0079132

10. Sommer M, Figueroa C, Kwauk C, Jones M, Fyles N. Attention to menstrual hygiene management in schools: An analysis of education policy documents in low- and middle-income countries. Int J Educ Dev [Internet]. 2017 Nov 1 [cited 2019 Apr 27];57:73–82. Available from: https://www.sciencedirect.com/science/article/pii/S0738059317302316?dgcid=raven_sd_recommender_email

11. Bott S, Jejeebhoy S, Shah I, Puri C. Towards adulthood: exploring the sexual and reproductive health of adolescents in South Asia [Internet]. 2003 [cited 2019 Apr 28]. Available from: https://apps.who.int/iris/bitstream/handle/10665/42781/9241562501.pdf;jsessionid=0C1BC887BCFF72D3EA7BD9118798089E?sequence=1

12. Jasper C, Le T-T, Bartram J, Jasper C, Le T-T, Bartram J. Water and Sanitation in Schools: A Systematic Review of the Health and Educational Outcomes. Int J Environ Res Public Health [Internet]. 2012 Aug 3 [cited 2019 Apr 28];9(8):2772–87. Available from: http://www.mdpi.com/1660-4601/9/8/2772

13. Kaur R, Kaur K, Kaur R. Menstrual Hygiene, Management, and Waste Disposal: Practices and Challenges Faced by Girls/Women of Developing Countries. J Environ Public Health [Internet]. 2018 Feb 20 [cited 2019 Apr 28];2018:1–9. Available from: https://www.hindawi.com/journals/jeph/2018/1730964/

14. Azage M, Ejigu T, Mulugeta Y. Menstrual hygiene management practices and associated factors among urban and rural adolescents in bahir dar city administration, northwest Ethiopia. Ethiop J Reprod Heal [Internet]. 2018 Dec 7 [cited 2019 Apr 13];10(4). Available from: http://ejrh.org/index.php/ejrh/article/view/207

15. von Elm E, Altman DG, Egger M, Pocock SJ, Gøtzsche PC, Vandenbroucke JP, et al. The Strengthening the Reporting of Observational Studies in Epidemiology (STROBE) statement: guidelines for reporting observational studies. [Internet]. Vol. 147, Annals of internal medicine. 2007 [cited 2019 Jul 13]. p. 573–7. Available from: http://www.ncbi.nlm.nih.gov/pubmed/17938396

16. Upashe SP, Tekelab T, Mekonnen J. Assessment of knowledge and practice of menstrual hygiene among high school girls in Western Ethiopia. BMC Women’s Health [Internet]. 2015 Dec 14 [cited 2019 Apr 5];15(1):84. Available from: http://bmcwomenshealth.biomedcentral.com/articles/10.1186/s12905-015-0245-7

17. Ketema Gultie T. Practice of Menstrual Hygiene and Associated Factors among Female Mehalmeda High School Students in Amhara Regional State, Ethiopia. Sci J Public Heal [Internet]. 2014 [cited 2019 Apr 8];2(3):189. Available from: http://www.sciencepublishinggroup.com/journal/paperinfo.aspx?journalid=251&doi=10.11648/j.sjph.20140203.18

18. Tegegne TK, Sisay MM. Menstrual hygiene management and school absenteeism among female adolescent students in Northeast Ethiopia. BMC Public Health [Internet]. 2014 Dec 29 [cited 2019 Apr 8];14(1):1118. Available from: https://bmcpublichealth.biomedcentral.com/articles/10.1186/1471-2458-14-1118

19. O. OM, S. UA, J. GG, T. AJ. Menstrual health: the unmet needs of adolescent girls in Sokoto, Nigeria. Sci Res Essays [Internet]. 2012 Jan 23 [cited 2019 Jul 10];7(3):410–8. Available from: https://academicjournals.org/journal/SRE/article-abstract/119BFB829292

20. Erulkar AS, Ferede A, Woldemariam WA, Unfpa G, Amdemikael H, Gebremedhin B, et al. Ethiopia Young adult SurvEY a StudY in SEvEn rEgionS Population Council [Internet]. 2010 [cited 2019 Jul 9]. Available from: www.popcouncil.org

21. Morrison J, Basnet M, Bhatta A, Khimbanjar S, Joshi D, Baral S. Menstrual hygiene management in Udaypur and Sindhuli districts of Nepal [Internet]. 2016 [cited 2019 Jul 9]. Available from: www.wateraid.org/mhm

